# Considering African savanna elephant conservation as a nature-based climate solution

**DOI:** 10.64898/2026.11.17.767383

**Authors:** Oswald J. Schmitz, Paradzayi Tagwireyi

## Abstract

The conservation of African savanna ecosystems is recognized as making important contributions toward capturing and storing atmospheric carbon in vegetation and in soils to help mitigate climate change. Also, savannas are important for elephant conservation, which is an increasing priority throughout Africa. However, there is concern that elephants can damage and transform vegetation thereby compromising savanna ecosystems’ capacity to capture and store carbon. Yet, elephants can have positive effects by releasing body wastes that contribute to nutrient cycling to enhance vegetation productivity (and hence carbon capture). The release of carbon in dung and trampling of soils can enhance long-term carbon storage in soils. Thus, an ecologically holistic assessment of net elephant effects on carbon capture and storage requires accounting for the negative and positive effects together. We report on a holistic assessment of elephant effects using a carbon cycle model that compares savanna ecosystem carbon capture and storage between scenarios with and without elephants. The modeling estimates that elephants at a density of ∼0.5 km^-2^ could increase average net ecosystem carbon capture by 1.4× (range of uncertainty of 0.4× – 5.5×). Elephants are estimated to increase average long-term aboveground carbon stocks by 6.5× (uncertainty of 2× - >10×). Average long-term soil carbon storage is estimated to increase by1.7× times (uncertainty of 0.6× – 5.3×). The estimated annual carbon capture is large enough to be considered a negative emission in national emissions accounting from land use change, suggesting that savanna elephant conservation should be considered as a nature-based climate solution.

## 1. Introduction

Two of the most important challenges confronting conservation scientists, policymakers, and practitioners is mitigating climate change and biodiversity loss (Dinnerstein et al. 2020, Seddon et al. 2021, Pörtner et al. 2023). Actions to address each challenge have tended to be treated separately. However, there is growing evidence that conserving biodiversity, especially animals, may help to mitigate climate change as part of nature-based climate solutions (Schmitz et al. 2023). Animals can contribute to nature-based climate solutions (NbCS) through their functional roles in ecosystems (Schmitz et al. 2014, 2018, Cromsigt et al. 2018, Malhi et al. 2022). These functional roles can ensure that the structure and functioning of ecosystems remains intact, thereby preventing carbon emissions (i.e., avoided emissions) that otherwise would result from the degradation of ecosystems. They can also enhance the capacity of ecosystems to capture and store additional carbon (i.e., enhanced sink strength). Thus, animal conservation aimed at restoring and conserving not just species but also their functional roles within ecosystems may offer the additional dividend of helping to arrest and mitigate climate change (Schmitz and Sylvén 2023).

An animal conservation strategy that deliberately addresses both challenges is known as animating the carbon cycle (Schmitz 2014). Animating the carbon cycle (ACC) recognizes that animals, particularly large vertebrates, can have important effects on ecosystem carbon cycling despite their smaller total biomass relative to other biotic entities (e.g., plants or microbes) that are important drivers of whole ecosystem carbon cycling (Schmitz et al. 2014; 2018). Empirical estimates from studies of terrestrial large animals indicate that they can, on average, help increase ecosystem carbon capture and storage by 1.2 × – 6 × above that in their absence (Schmitz and Leroux 2026). This high potential of ACC as an NbCS makes it an appealing way to promote wildlife conservation everywhere. However, depending on ecosystem type, animal species may reduce carbon capture and storage by 0.5× – 3× (Schmitz and Leroux 2020). We are thus at a juncture where careful accounting of species functional effects on ecosystem carbon budgets is necessary if conservation aims to strive for reliable and ecologically accurate biodiversity recovery and conservation using the concept of ACC. This issue is especially germane when considering the functional role of large mammals such as elephants in African ecosystems (Burak et al., 2024; Duvall et al., 2024; Abraham et al., 2026).

Opinion is divided about whether African savanna elephants (*Loxodonta africana*) have a beneficial effect in savanna-woodland ecosystems. Indeed, they have long been considered to be dominant agents of vegetation destruction (Ben-Shahar, 1993; Midgley et al., 2005; Shannon et al., 2011; Asner and Levick, 2012; Owen-Smith et al.; 2019; Chakuya et al., 2022; Mukomberanwa et al., 2024), which could modify vegetation community structure, and lead to loss of plant and animal biodiversity across savanna-woodland landscapes (Cumming et al., 1997; Holdo, 2007; Gandiwa et al., 2011; Valeix et al., 2011; Zyambo, 2016; Shannon et al., 2011; Guldemond et al., 2017; Davies et al., 2018; O’Connor et al., 2024). Yet more recently their impacts on vegetation are also considered neutral or beneficial for maintaining biodiversity (Skarpe et al., 2004; Guldemond et al., 2017; Gordon et al., 2023; Western and Mose, 2023; Gijsman et al., 2026; Kuiper et al., 2026), plant community stability (Jeltsch et al., 2000; Baxter and Getz, 2005, Ferry et al., 2021; Thel et al., 2025), and community resilience to the effects of climate change (Stevens et al., 2016; Gordon et al., 2022; Simba et al., 2024).

Recent evidence from studies evaluating impacts by functionally similar African forest elephant’s (*L. cyclotis*) (Poulson et al., 2018; Berzaghi et al. 2019; Gupta et al. 2026), would suggest that savanna elephants could contribute to maintain climate resilience by enhancing the capacity of savanna-woodland ecosystems to capture and store carbon. But here too opinion is divided. Losses of overstory canopy tree and understory shrub and grass diversity and biomass due to savanna elephant foraging, trampling and tree felling leads to conclusions that elephants may limit or even reduce the ecosystems’ capacity to store carbon (Pellegrini et al., 2017; Davies and Asner, 2019; Wigley et al., 2020; Wessels et al., 2026). Yet analyses of the carbon released to soils in the form of dung and decaying wood leads to conclusions that savanna elephants may promote carbon storage (Sandhage-Hofmann et al., 2021; 2022; Kahumba and Tefera, 2024; Kinderman et al., 2025). Furthermore, elephant carcasses could contribute substantial amounts of carbon to localized soil pools, but nutrients released from those carcasses could also stimulate soil respiration, leading to soil carbon loss (Reed et al. 2025).

Reconciling these different conclusions requires overcoming the largely piecemeal approach currently taken by ecological science, in which studies focus on one or two components of the entire carbon cycle. An ecologically holistic assessment of elephant effects via ACC explicitly requires accounting for the negative and positive effects together to estimate elephant *net effects* on ecosystem carbon capture and storage (Schmitz et al. 2018). Thus, ACC requires broadening consideration beyond elephant damage to plant carbon and elephant enhancement of soil carbon to include six kinds of effects that impact both carbon storage (i.e. stocks) and carbon uptake and releases (i.e., fluxes) among ecosystems components (i.e., soils, plants and animals), and between ecosystem components and the atmosphere. These effects include (Fig 1): (i) elephant foraging and plant biomass carbon loss; (ii) elephant foraging effects on plant atmospheric carbon uptake and plant carbon release in respiration (i.e., plant net primary productivity [NPP], aka net ecosystem carbon capture); (iii) elephant recycling release of nutrients in urine and dung that could stimulate NPP; (iv) elephant contributions to soil carbon storage via dung and carcass deposition; (v) elephant alteration of soil respiratory CO_2_ release via trampling effects; and (vi) elephant respiratory CO_2_ and enteric methane release. Moreover, quantification of (ii), (v) and (vi) together is needed to estimate another important flux, variously called net ecosystem exchange (NEE) or net ecosystem carbon balance (NECB), which essentially quantifies the net capacity of the whole ecosystem to capture carbon, aka, ecosystem carbon sink strength (Chapin et al., 2006; Rizzuto et al. 2024). Such a full accounting has yet to be done empirically for elephants in any savanna ecosystem worldwide (Hyvarinen et al. 2021). This leaves considerable uncertainty about whether African savanna elephant conservation can be considered as a nature-based climate solution.

**Fig. 1.**
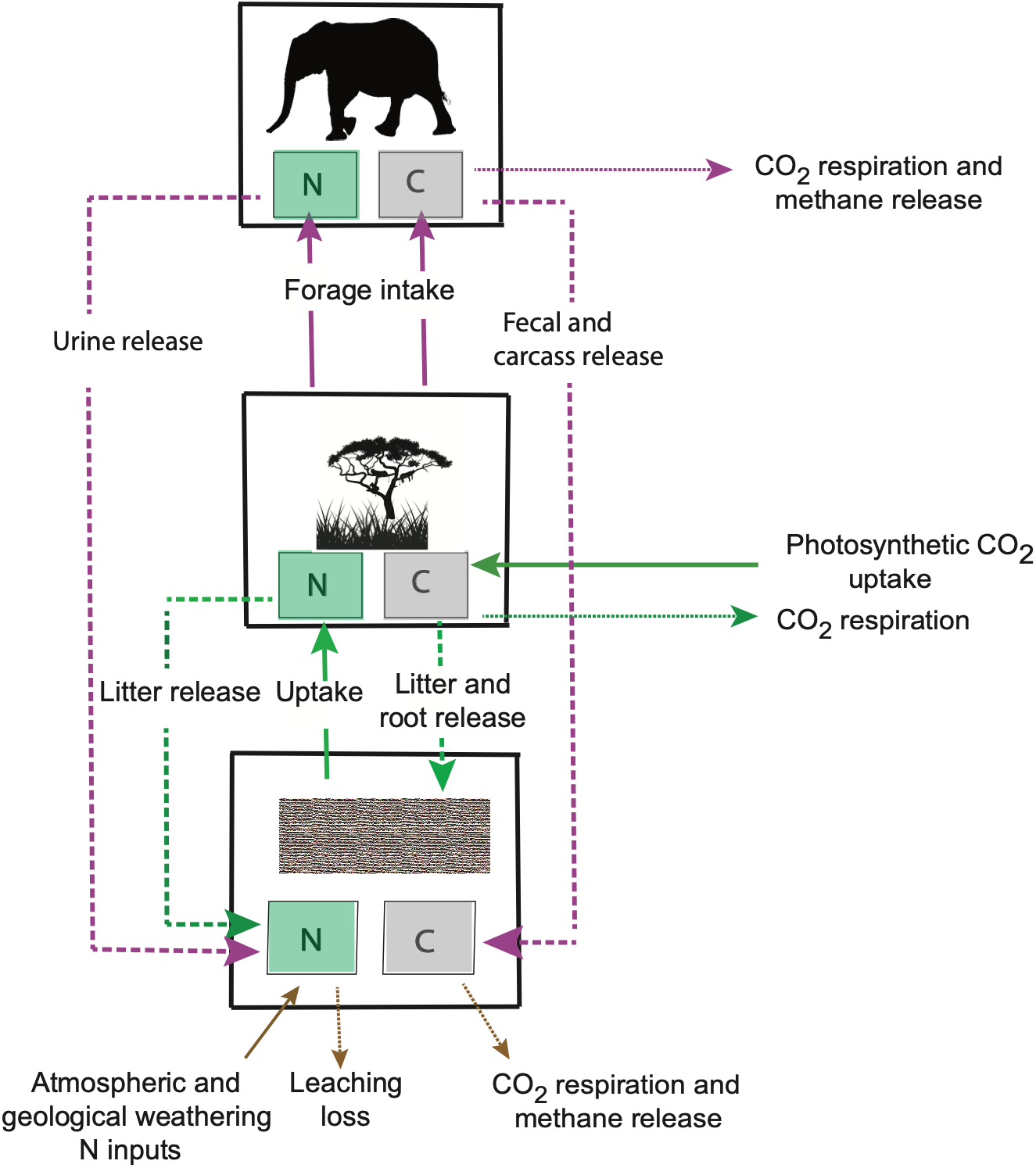
The processes involved in the uptake, release and storage of carbon (C) and nutrients (N) within a savanna-woodland ecosystem. The figure shows the pathways of C and N uptake between soil and vegetation, and vegetation and elephant compartments of the ecosystem and nutrient and carbon recycling feedbacks from vegetation and elephants to the soil compartment. The boxes represent C and N stocks and the lines represent fluxes. Solid lines represent direct uptake; dashed lines represent recycling feedback within the ecosystem; and dotted lines represent release from the ecosystem. The colors of the arrows indicate which ecosystem compartment is taking up or releasing C and N with magenta representing elephants, green representing vegetation and brown representing soil.

Addressing uncertainty about the impacts of large, widely roaming animals ideally involves obtaining robust measures of carbon stocks and fluxes between locations with similar biophysical ecosystem conditions, but where focal animals are present vs naturally absent or experimentally excluded (Schmitz et al., 2018; Schmitz and Leroux, 2026). Conducting such long-term experiments with elephants is logistically challenging across broad landscapes and is thus not widely done. When opportunities to leverage such enclosure experiments have arisen (e.g., Pellegrini et al., 2017; Wigley et al., 2020; Wessels et al., 2026), they provide important insights about elephant effects. Nonetheless, so far, they only measure one or a few of the six effects, which precludes a comprehensive understanding of elephant net effects.

Ecosystem modelling offers an alternative and immediate way to begin developing the kind of comprehensive understanding needed to address uncertainty (Schmitz and Leroux 2026). When parametrized using data from published studies, ecosystem models can be used to systematically explore scenarios that estimate and compare ecosystem carbon budgets in the presence and absence of focal animals. Here we report on an application of such modelling to examine elephant impacts on carbon capture and storage in savanna-woodland ecosystem plants and soils. Recognizing that no single model is likely to explain the entire variety of functions and patterns observed among savanna locations (Scholes and Archer, 1997; Jeltsch et al., 2000), our analysis nonetheless provides preliminary insight into how much carbon capture and storage could be attributed to the functional role of savanna elephants. We tailored a general animal-driven carbon cycle model (Rizzuto et al., 2024) to embody the salient principles of animal-driven savanna-woodland ecosystem carbon and nutrient dynamics (Skarpe et al., 2004; Holdo et al., 2007, du Toit et al., 2014, Coetsee et al., 2019; Huntley, 2023). The modeling reveals that savanna elephant conservation should be considered further as a feasible NbCS given that the magnitude of elephant effects are large enough to help reduce carbon emissions from land use and land use change. However, there are uncertainties related to parameter values and model estimates that need to be addressed with further empirical research to obtain accurate accounts of total carbon capture and storage. To this end, the modelling identifies crucial new field research that is needed to reduce uncertainty about elephant conservation as an NbCS.

## 2.0 Modeling elephant effects on savanna-woodland ecosystem carbon cycling

### 2.1 Focal savanna-woodland ecosystem type

Savanna elephants frequently occur in three broad types of savanna-woodland ecosystems: the wetter miombo, and the drier mopane and bushveld (Gobush et al. 2021). We focus our modeling on the mopane savannah-woodland. Mopane savanna-woodlands are widespread and dominant in arid, low-lying areas across much of Southern Africa, including northern Botswana, Namibia, Mozambique, northern South Africa, and Zimbabwe (Shorroks and Bates, 2015; Ossifo et al., 2026). Mopane tree leaves and twigs in addition to grasses are the predominant food source for savanna elephants (Kos et al., 2012; Sach et al. 2019). Moreover, elephant functional roles have been studied more extensively in this ecosystem type than in the other savanna-woodland types. Thus, studies in this ecosystem type provide the best body of scientific insights and data to date on aboveground vegetation biomass carbon and on soil carbon and elephant impacts, which can be used to assess the reliability of the model results.

### 2.2. Overview of the modeling approach

The modeling accounts for the flow and storage of carbon among soil, vegetation, and elephant ecosystem compartments and exchanges between the ecosystem and the atmosphere (Fig 1). The mathematical model describing the various process depicted in Fig 1 is presented in Table 1.

**Table 1.**
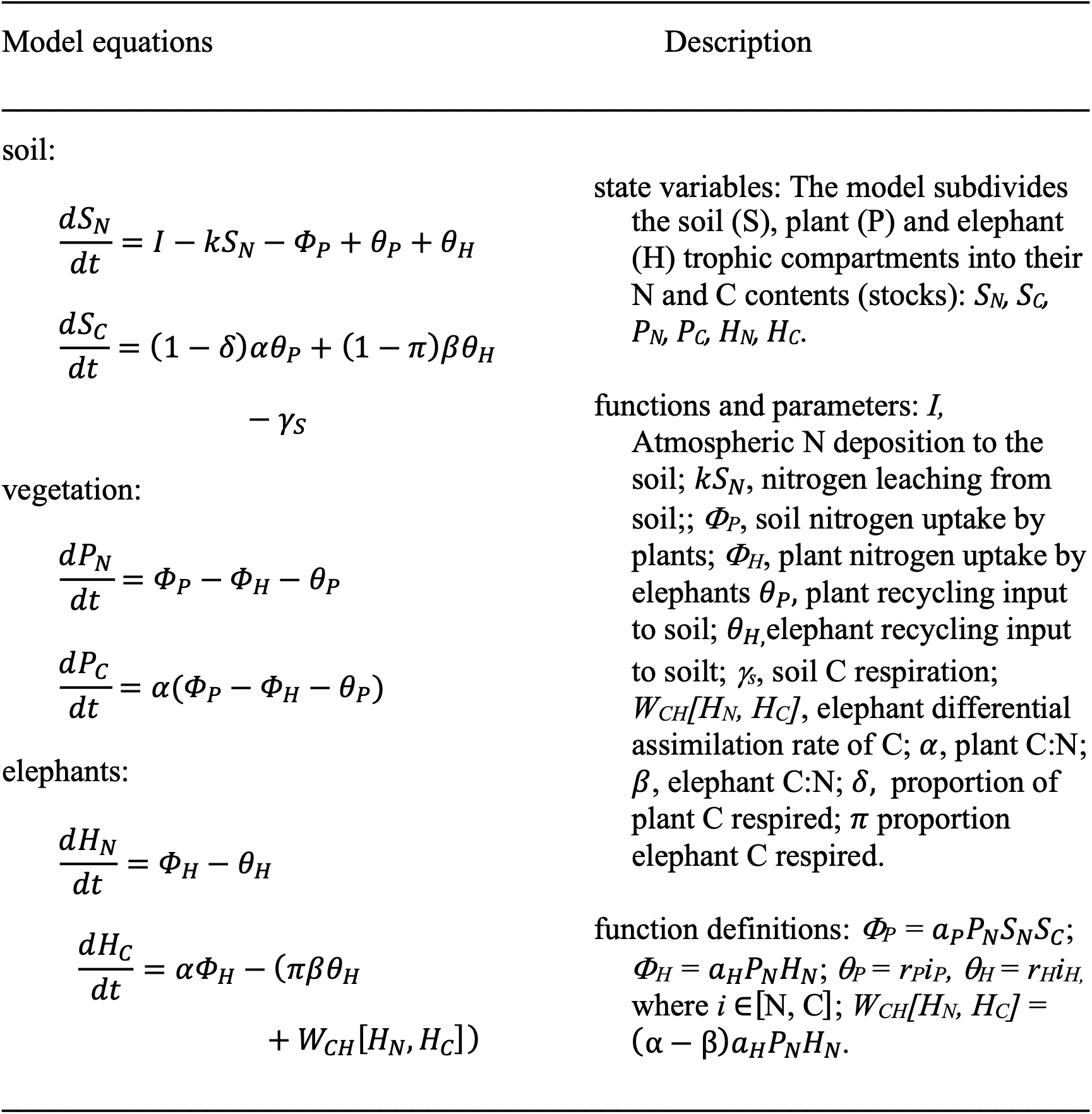
Mathematical equations used to characterize the processes of carbon and nitrogen uptake, flow and storage among soil, vegetation and elephant compartments of a Mopane ecosystem. The table provides variable, function and parameter definitions. See text Figure 1 for a conceptual diagram of the ecosystem model.

| Model equations | Description |
|---|---|
| soil: |  |
| $\frac{dS_N}{dt} = I - kS_N - \Phi_P + \theta_P + \theta_H$ | state variables: The model subdivides the soil (S), plant (P) and elephant (H) trophic compartments into their N and C contents (stocks): *S<sub>N</sub>, S<sub>C</sub>, P<sub>N</sub>, P<sub>C</sub>, H<sub>N</sub>, H<sub>C</sub>*. |
| $\frac{dS_C}{dt} = (1 - \delta)\alpha\theta_P + (1 - \pi)\beta\theta_H - \gamma_S$ | functions and parameters: *I,* Atmospheric N deposition to the soil; *kS<sub>N</sub>*, nitrogen leaching from soil;; *Φ<sub>P</sub>*, soil nitrogen uptake by plants; *Φ<sub>H</sub>*, plant nitrogen uptake by elephants *θ<sub>P</sub>*, plant recycling input to soil; *θ<sub>H,</sub>*elephant recycling input to soilt; *γ<sub>s</sub>*, soil C respiration; *W<sub>CH</sub>[H<sub>N</sub>, H<sub>C</sub>]*, elephant differential assimilation rate of C; *α*, plant C:N; *β*, elephant C:N; *δ,* proportion of plant C respired; *π* proportion elephant C respired. |
| vegetation: |  |
| $\frac{dP_N}{dt} = \Phi_P - \Phi_H - \theta_P$ |  |
| $\frac{dP_C}{dt} = \alpha(\Phi_P - \Phi_H - \theta_P)$ |  |
| elephants: |  |
| $\frac{dH_N}{dt} = \Phi_H - \theta_H$ |  |
| $\frac{dH_C}{dt} = \alpha\Phi_H - (\pi\beta\theta_H + W_{CH}[H_N, H_C])$ | function definitions: *Φ<sub>P</sub>* = *a<sub>P</sub>P<sub>N</sub>S<sub>N</sub>S<sub>C</sub>*; *Φ<sub>H</sub>* = *a<sub>H</sub>P<sub>N</sub>H<sub>N</sub>*; *θ<sub>P</sub>* = *r<sub>P</sub>i<sub>P</sub>,* *θ<sub>H</sub>* = *r<sub>H</sub>i<sub>H,</sub>* where *i* ∈[N, C]; *W<sub>CH</sub>[H<sub>N</sub>, H<sub>C</sub>]* = (α − β)*a<sub>H</sub>P<sub>N</sub>H<sub>N</sub>*. |

The full model (Table 1) accounts for ecosystem inputs including atmospheric N deposition to the ecosystem (I) and atmospheric C uptake by vegetation (*αΦ_P_*). The model accounts for storage via the redistribution of C and N among ecosystem compartments through: (i) vegetation uptake of soil nitrogen N (*Φ_P_*), (ii) uptake of vegetation C (*αΦ_H_*) and N (*Φ_H_*) by elephant herbivory, and recycling of plant organic matter to soils (iii) via plant litter (i.e., vegetation C (*(1 – δ)αθ_P_*) and N (*θ_P_*)) and (iv) via elephant waste (i.e., C ((*1 – π*)*βθ_H_*) and N (*θ_H_*)) which includes feces, urine, and body carcasses. Modeled ecosystem outputs include soil N leaching out of the ecosystems (*K*), and respiratory C release to the atmosphere from soils (*γ_S_*), vegetation (*δαθ_P_*), and elephants (*πβθ_H_ + W_CH_*[*H_N_*, *H_C_*]). The term *W_CH_*[*H_N_*, *H_C_*] partitions carbon intake into respiration versus recycling, reflecting herbivore stoichiometric homeostasis i.e., the need to maintain a fixed C:N ratio (*β*) despite consuming plant material with a typically higher C/N (*α*) (Rizzuto et al. 2024). Generally, plant biomass has high C and low N whereas herbivore biomass has low C high and high N, hence herbivore nutrient limitation (McNaughton et al. 1988, Holdo et al. 2007). Hence, the modeling accounts for the rate limiting influence of essential nutrients on carbon cycling (Rastetter 2011; Rizzuto et al. 2024; Zaehle et al. 2014). The focus on C and N stems from their known importance in savanna biogeochemical cycling (McNaughton et al., 1988; Coetsee et al., 2019; Huntley, 2023). The mathematical model is solved for steady state conditions that obey fundamental mass balance requirements in which ecosystem C and N inputs equal outputs plus storage within ecosystem compartments, i.e., it produces a balanced ecosystem carbon budget (Rizzuto et al. 2024). The steady state model solutions (see Appendix A: A1) are used to estimate elephant effects on the net amount of carbon that is captured by plants (NPP), the carbon sink strength of the ecosystem (NECB) and the amount of carbon stored in soil and in plant biomass.

The carbon estimates were produced by parametrizing the model with empirical data. Ideally, the full suite of empirical data to parameterize the model would come from a detailed empirical study of the interaction of elephants, trees and soil within the focal location. However, such a study does not exist. Thus, empirical data to estimate parameters were obtained from the published literature for mopane ecosystems throughout Africa. A full description of the parameter calculations, including assumptions and the sources of the data, is presented in Appendix A: A2.

The parameters were estimated to account for elephant effects for an entire year that includes a 181-day wet season in which vegetation is productive and captures atmospheric carbon, and a 184-day dry season in which plant carbon capture is considerably reduced if not stopped due to plant dormancy, but where plant and soil CO_2_ respiration, and elephant foraging, CO_2_ respiration and methane release continues. Thus, dry season effects on NECB and plant standing carbon were added or debited from the wet season values when calculating a yearly net carbon budget.

Using the parameter values (Table 2) as starting points, the model was run for two scenarios: (1) carbon uptake and storage in (1) the absence of elephants (No Elephants); and (2) the presence of elephants (With elephants). These scenarios facilitated estimating the amount of additional carbon that could be gained by conserving elephants in the ecosystem—i.e., the ACC potential of conserving elephants. ACC potential represents net effect of elephants on the savanna ecosystem carbon dynamics and is quantified as the difference between the two model scenarios.

**Table 2.**
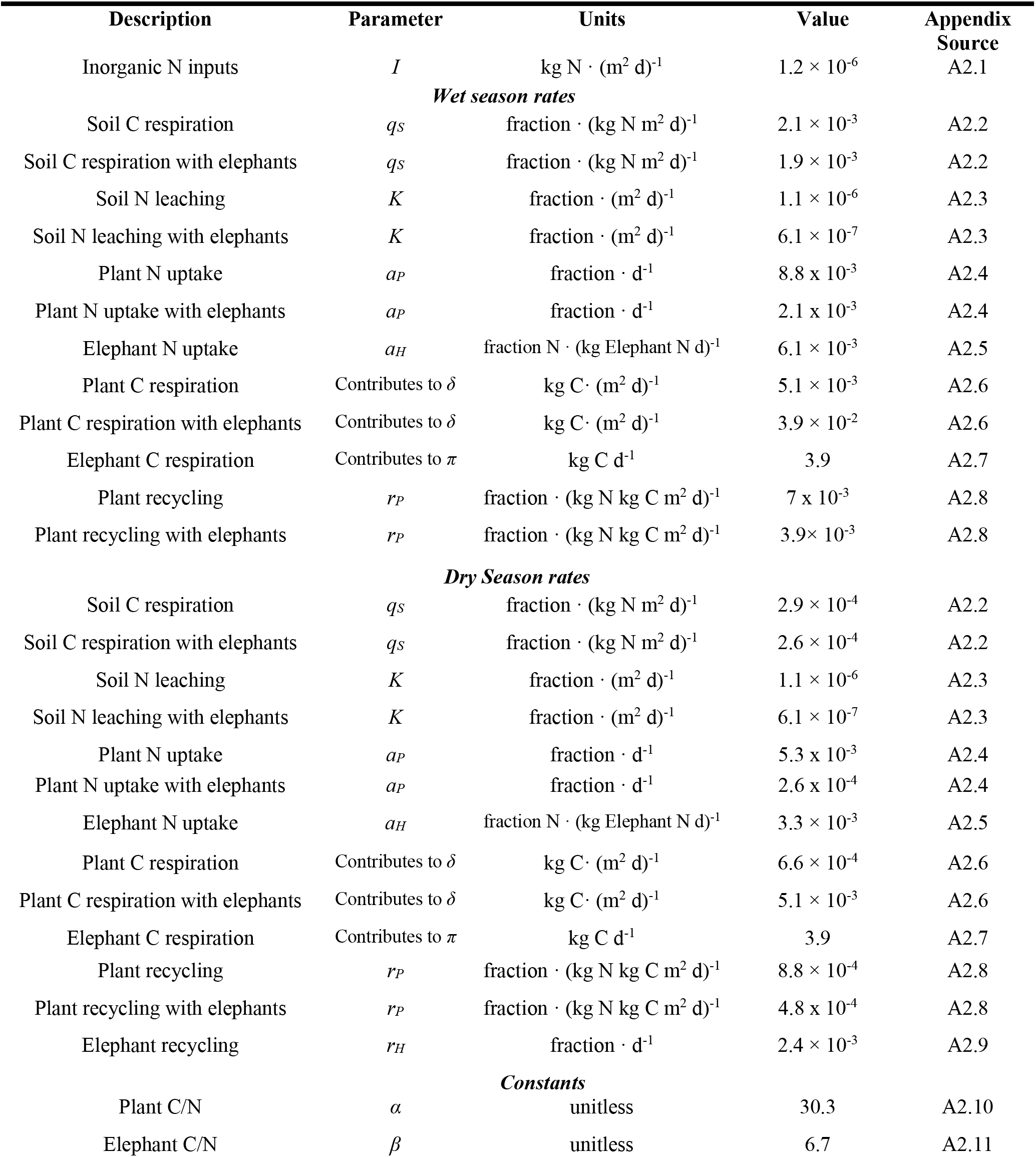
Parameters, their units and average empirical values obtained from the literature to numerically analyze savanna elephant effects on carbon capture and storage using steady state solutions of the dynamical systems model presented in Table 1.

| **Description** | **Parameter** | **Units** | **Value** | **Appendix Source** |
|---|---|---|---|---|
| Inorganic N inputs | *I* | kg N · (m<sup>2</sup> d)<sup>-1</sup> | 1.2 × 10<sup>-6</sup> | A2.1 |
| ***Wet season rates*** |  |  |  |  |
| Soil C respiration | *q<sub>S</sub>* | fraction · (kg N m<sup>2</sup> d)<sup>-1</sup> | 2.1 × 10<sup>-3</sup> | A2.2 |
| Soil C respiration with elephants | *q<sub>S</sub>* | fraction · (kg N m<sup>2</sup> d)<sup>-1</sup> | 1.9 × 10<sup>-3</sup> | A2.2 |
| Soil N leaching | *K* | fraction · (m<sup>2</sup> d)<sup>-1</sup> | 1.1 × 10<sup>-6</sup> | A2.3 |
| Soil N leaching with elephants | *K* | fraction · (m<sup>2</sup> d)<sup>-1</sup> | 6.1 × 10<sup>-7</sup> | A2.3 |
| Plant N uptake | *a<sub>P</sub>* | fraction · d<sup>-1</sup> | 8.8 x 10<sup>-3</sup> | A2.4 |
| Plant N uptake with elephants | *a<sub>P</sub>* | fraction · d<sup>-1</sup> | 2.1 x 10<sup>-3</sup> | A2.4 |
| Elephant N uptake | *a<sub>H</sub>* | fraction N · (kg Elephant N d)<sup>-1</sup> | 6.1 × 10<sup>-3</sup> | A2.5 |
| Plant C respiration | Contributes to *δ* | kg C· (m<sup>2</sup> d)<sup>-1</sup> | 5.1 × 10<sup>-3</sup> | A2.6 |
| Plant C respiration with elephants | Contributes to *δ* | kg C· (m<sup>2</sup> d)<sup>-1</sup> | 3.9 × 10<sup>-2</sup> | A2.6 |
| Elephant C respiration | Contributes to *π* | kg C d<sup>-1</sup> | 3.9 | A2.7 |
| Plant recycling | *r<sub>P</sub>* | fraction · (kg N kg C m<sup>2</sup> d)<sup>-1</sup> | 7 x 10<sup>-3</sup> | A2.8 |
| Plant recycling with elephants | *r<sub>P</sub>* | fraction · (kg N kg C m<sup>2</sup> d)<sup>-1</sup> | 3.9× 10<sup>-3</sup> | A2.8 |
| ***Dry Season rates*** |  |  |  |  |
| Soil C respiration | *q<sub>S</sub>* | fraction · (kg N m<sup>2</sup> d)<sup>-1</sup> | 2.9 × 10<sup>-4</sup> | A2.2 |
| Soil C respiration with elephants | *q<sub>S</sub>* | fraction · (kg N m<sup>2</sup> d)<sup>-1</sup> | 2.6 × 10<sup>-4</sup> | A2.2 |
| Soil N leaching | *K* | fraction · (m<sup>2</sup> d)<sup>-1</sup> | 1.1 × 10<sup>-6</sup> | A2.3 |
| Soil N leaching with elephants | *K* | fraction · (m<sup>2</sup> d)<sup>-1</sup> | 6.1 × 10<sup>-7</sup> | A2.3 |
| Plant N uptake | *a<sub>P</sub>* | fraction · d<sup>-1</sup> | 5.3 x 10<sup>-3</sup> | A2.4 |
| Plant N uptake with elephants | *a<sub>P</sub>* | fraction · d<sup>-1</sup> | 2.6 x 10<sup>-4</sup> | A2.4 |
| Elephant N uptake | *a<sub>H</sub>* | fraction N · (kg Elephant N d)<sup>-1</sup> | 3.3 × 10<sup>-3</sup> | A2.5 |
| Plant C respiration | Contributes to *δ* | kg C· (m<sup>2</sup> d)<sup>-1</sup> | 6.6 × 10<sup>-4</sup> | A2.6 |
| Plant C respiration with elephants | Contributes to *δ* | kg C· (m<sup>2</sup> d)<sup>-1</sup> | 5.1 × 10<sup>-3</sup> | A2.6 |
| Elephant C respiration | Contributes to *π* | kg C d<sup>-1</sup> | 3.9 | A2.7 |
| Plant recycling | *r<sub>P</sub>* | fraction · (kg N kg C m<sup>2</sup> d)<sup>-1</sup> | 8.8 × 10<sup>-4</sup> | A2.8 |
| Plant recycling with elephants | *r<sub>P</sub>* | fraction · (kg N kg C m<sup>2</sup> d)<sup>-1</sup> | 4.8 x 10<sup>-4</sup> | A2.8 |
| Elephant recycling | *r<sub>H</sub>* | fraction · d<sup>-1</sup> | 2.4 × 10<sup>-3</sup> | A2.9 |
| ***Constants*** |  |  |  |  |
| Plant C/N | *α* | unitless | 30.3 | A2.10 |
| Elephant C/N | *β* | unitless | 6.7 | A2.11 |

The parameter values were also adjusted individually to explore the sensitivity of model estimates to changes in each of the parameters (i.e., a sensitivity analysis). The model parameters were further changed singly and in various combinations of one-another to examine the uncertainty around estimates of carbon capture and storage for each scenario (i.e., an uncertainty analysis).

The modeling assessed the potential effects of elephants on carbon capture and storage at an elephant density of approximately 0.5-0.7 individuals per km^2^. This density falls in the mid-range of natural and managed densities throughout much of the savanna elephants’ range (Guldemond et al., 2017) and is considered an acceptable density from a conservation and management perspective (Scheiter and Higgins, 2012; Guldemond et al., 2017; Kuiper et al., 2026). The modeling analyses provide *a first approximation estimate* of the potential impacts that elephants have on the Mopane ecosystem carbon capture and storage. The model estimates are compared with field measurements of Mopane savanna ecosystem carbon reported in the literature to assess the reliability of the model estimates.

### 2.3 Model implementation

Each modeling scenario (i.e., animal absent; animal present) has one equilibrium where all state variables (soil = *S_i_,* plant = *P_i_,* herbivore *= H_i_*; where *i* identifies the stock as either carbon (C) or nitrogen (N)) are positive, and thus biologically meaningful (see Appendix A: A1). Appendix A1 also includes equations used to estimate NPP and NECB. The stock (soil and plant C) and flux (NPP, NECB) estimates were derived by substituting model parameters (Table 2) into the respective equations. The No elephant scenario was simulated by setting herbivore consumption and waste recycling parameters to zero i.e., *a_H_* and *r_H_* = 0 in *Φ_H_*, *θ_H_*. The With elephant scenario was simulated by including values for the herbivore consumption and recycling parameters (Table 2). We also adjusted other parameters to account for elephant effects, i.e., elephant trampling on soil C respiration and N leaching; plant N uptake and C respiration due to plant stress from herbivore foraging impacts; and plant litter recycling due to reduced available plant biomass consequent to foraging (Table 2; Appendix A: A2). Parameter values in Table 2 were entered into a structured Excel spreadsheet designed to automatically populate model equations for equilibrium soil and plant C, and NPP and NECB. All estimates were made in units of kg C, m^2^ and day to facilitate comparisons with published field measurements.

#### 2.2.1 Estimates of carbon capture and storage for the No elephant scenario

The No elephant scenario was used to obtain baseline estimates of carbon capture and storage in plant and soil trophic compartments of the ecosystem and associated NPP and NECB. The model estimates total plant carbon does not distinguish between carbon in aboveground biomass (stems and leaves) and in belowground biomass (roots). However, field studies available to validate the model estimates report measures in terms of aboveground biomass carbon only. The model estimates of total plant carbon were thus adjusted downward given that plant biomass in mopane savanna-woodland can be comprised of 14% - 21% roots (Lisboa et al. 2025).

We conducted a sensitivity analysis by varying the seven key parameters: (i) atmospheric deposition of inorganic N, (ii) soil C leaching rate, (iii) soil N leaching rate, (iv) plant N uptake rate, and (v) plant C respiration rate; (vi) plant litter recycling rate; and (vii) plant C/N. We varied parameter values by up to +25% and -25% of the mean, based on insights from a sensitivity analysis of the general model (Rizutto et al. 2024), which indicates that a ± 25% range will produce robust estimates of uncertainty. The parameters were varied one-at-a time to assess sensitivity. A change in model estimate > 25% was considered *sensitive*, a change = 25% was considered *equal*, and a change < 25% was considered *insensitive*).

The degree of uncertainty in model estimates (uncertainty_C-estimates_) was ascertained by solving for multiple balanced carbon budgets. This was done by adjusting each parameter singly and in combination to generated estimates of carbon capture and storage. The parameters were then reset to their original values before exploring new combinations. The parameters were adjusted by up to ± 50% of that reported in Table 2, which encompasses the magnitudes of calculated ranges in parameter values obtained from the literature (Appendix A: A2).

#### 2.2.2 Estimates of carbon capture and storage for the With elephant scenario

Elephant effects on carbon dynamics were estimated by adjusting the seven key parameters used in the modeling of No elephant plant-soil dynamics to account for the elephant effects on biogeochemical processes (Table 2). The rationale for those parameter adjustments is presented in Appendix A: A2. As well, the modeling scenario used additional parameters to characterize elephant foraging, recycling, respiration and body C/N.

We estimated net carbon capture (NPP and NECB) and soil and plant carbon stock such that the estimated elephant density approximated 0.5 - 0.7 animals km^-2^. Elephant density was estimated in several steps. We first converted the carbon stock estimates of the elephant trophic compartment into whole-animal aggregate density (individuals km^-2^) by dividing the estimated elephant carbon stock (kg C km^-2^) by the product of a weighted average elephant population body mass and % carbon in elephant body mass (see Appendix A - A2.5). In estimating NECB, we also accounted for enteric methane (CH_4_) emissions from elephants. As no empirical measurement of elephant CH_4_ emission measurements exist yet, we used an allometric relationship presented in Smith et al. (2015) to estimate CH_4_ emission per average-sized elephant individual and scaled that to density.

We assessed model sensitivity to changes in the seven key plant-soil parameters impacted by elephant effects, and four additional elephant-related parameters: (i) elephant N uptake rate; (ii) elephant C respiration; (iii) elephant recycling; and (iv) elephant C/N. The sensitivity analysis involved varying each parameter one at a time by ± 25% and ascertaining how much the model estimate of plant and soil C, and NPP and NECB changed. Uncertainty_C-estimates_ was again estimated by solving for multiple balanced carbon budgets. The parameters were reset to their original values before exploring new combinations

The model estimates of plant carbon were again adjusted downward by 14% - 21% of total plant biomass to account for root biomass. Furthermore, elephants are also known to reduce aboveground plant biomass by felling trees. Such effects can reduce standing live aboveground biomass by 25% - 30% (Chakuya et al., 2022, Kindermann et al., 2022, Coetsee et al., 2023, O’Connor et al., 2024; Archibald et al., 2026). Thus, in addition to applying the correction for estimating aboveground plant biomass carbon, estimates were further reduced by 30% to account for additional elephant damages not represented in measures of biomass forage consumption rates.

#### 2.2.3 Estimates of the contribution of elephants to ecosystem carbon dynamics

The average change in estimated amount of carbon capture and storage due to elephant presence was calculated as the ratio of With elephant mean soil and plant C stocks, and NPP and NECB to the respective No elephant mean estimates. However, due to uncertainty about each mean estimate (uncertainty_C-estimate_) we also calculated uncertainty ranges for changes due to elephant effects (uncertainty_EE_).

The uncertainty_EE_ range was calculated using two endpoints. The first range endpoint was calculated as the ratio of the maximum uncertainty_C-estimate_ range value for the With elephant estimate of carbon capture or storage and the minimum uncertainty_C-estimate_ range value for respective estimate for the No elephant scenario. The second endpoint was calculated as the ratio of the minimum uncertainty_C-estimate_ range value for the With elephant estimate and the maximum uncertainty_C-estimate_ range value for the No elephant scenario. The uncertainty_EE_ ranges could alternatively be calculated as the ratios of maximum to maximum and minimum to minimum uncertainty_C-estimate_ range values between With and No elephant scenarios. This approach would produce narrower uncertainty_EE_ ranges. However, such calculations require assuming that environmental conditions are consistent between With and No elephant scenarios. This assumption may not be met because elephant presence can alter local biophysical conditions and plant recycling processes through trampling that compacts the soil and reduces soil respiration. Thus, the approach we used to calculate uncertainty_EE_ accounts for the potential for differences in biophysical conditions to occur between scenarios.

## 3.0 Model estimates of carbon capture and storage

The estimated soil and plant carbon stocks and NPP and NECB fluxes are presented in Table 3 along with measurements reported in the published literature. The reported values are ranges determined from the model uncertainty_C-estimate_ analysis or range of empirical values reported in the literature. The model estimates are on the same order of magnitude as empirical values, although the high end of the model estimated uncertainty range exceeds that for the measured range for soil C stocks in both scenarios, and for plant C stocks in the With elephant scenario (Table 3). Nevertheless, the comparability of the estimated and measure ranges provides some confidence in the reliability of the model estimates.

**Table 3.** Modeled and empirical estimates of mopane savanna ecosystem carbon storage (stocks) in soil and plants, and ecosystem carbon exchange (fluxes) as plant net CO_2_ uptake (NPP), and net ecosystem carbon sink strength (aka net ecosystem carbon balance; NECB). The reported values are ranges between minimum and maximum values estimated by the model uncertainty analysis (Modeled) and reported in the literature (Empirical). Model predictions are for steady state elephant densities of 0.70 ± 0.15 individuals per km^2^.

| Variable | Modeled | Empirical | Source |
|---|---|---|---|
| **No Elephant** |  |  |  |
| ***stocks*** |  |  |  |
| soil C (kg C m<sup>-2</sup>) | 0.35 – 1.20 | 0.3 – 0.65 | 1 |
| aboveground plant C (kg C m<sup>-2</sup>) | 0.33 – 1.11 | 0.1 – 3.4 | 2,3 |
| ***fluxes*** |  |  |  |
| *wet season* |  |  |  |
| NPP (kg C m<sup>-2</sup> d<sup>-1</sup>) | 1.3 x 10<sup>-3</sup> – 4.9 x 10<sup>-3</sup> | 1.0 x 10<sup>-3</sup> – 4 x 10<sup>-3</sup> | 4,5 |
| NECB (kg C m<sup>-2</sup> d<sup>-1</sup>) | 1.3 x 10<sup>-3</sup> – 4.9 x 10<sup>-3</sup> | 1.0 x 10<sup>-3</sup> – 5.1 x 10<sup>-3</sup> | 5,6 |
| *dry season* |  |  |  |
| NPP (kg C m<sup>-2</sup> d<sup>-1</sup>) | 3.6 x 10<sup>-5</sup> – 4.8 x10<sup>-4</sup> | 0 – 9 x 10<sup>-4</sup> | 4,5 |
| NECB (kg C m<sup>-2</sup> d<sup>-1</sup>) | 3.6 x 10<sup>-5</sup> – 4.8 x10<sup>-4</sup> | 0 - 8 x 10<sup>-4</sup> | 5 |
| **With Elephant** |  |  |  |
| ***stocks*** |  |  |  |
| soil C (kg C m<sup>-2</sup>) | 0.72 – 1.86 | 0.15 – 1.5 | 7,8,9 |
| aboveground plant C (kg C m<sup>-2</sup>) | 2.33 – 7.07<sup>*</sup> | 0.32 – 6 | 3<sup>**</sup>,7,10 |
| ***fluxes*** |  |  |  |
| *wet season* |  |  |  |
| NPP (kg C m<sup>-2</sup> d<sup>-1</sup>) | 1.9 x 10<sup>-3</sup> – 6.5 x 10<sup>-3</sup> | NA |  |
| NECB (kg C m<sup>-2</sup> d<sup>-1</sup>) | 1.9 x 10<sup>-3</sup> – 6.5 x 10<sup>-3</sup> | N/A |  |
| *dry season* |  |  |  |
| NPP (kg C m<sup>-2</sup> d<sup>-1</sup>) | 2.9 x 10<sup>-4</sup> – 8.7 x10<sup>-4</sup> | N/A |  |
| NECB (kg C m<sup>-2</sup> d<sup>-1</sup>) | 2.9 x 10<sup>-4</sup> – 8.7 x10<sup>-4</sup> | N/A |  |

1. Bird et al. 2004; 2. Macôo et al. 2025 and Ossifo et al. 2026; 3. Wessells et al. 2026; 4. Veenendaal et al 2004; 5. Jin et al. 2013; 6. Kutsch et al. 2008, 7. Kinderman et al. 2025; 8. Kahumba and Tefera 2024; 9. Cambule et al. 2014 in locations where elephants historically occur (cf. Roque et al 2022), 10. Ben-Shahar 1996.
N/A = not available
<sup>\*</sup> Assumes an average 30% loss of biomass due to elephant felling
<sup>\*\*</sup> Values used are for elephant density < 1 km<sup>2</sup>

The model estimates are variously sensitive to changes in parameter values (Fig. 2). In the No elephant scenario, model estimates of carbon stocks and fluxes were most sensitive to changes in soil N leaching rate and plant N uptake rate, and least sensitive to changes in plant C respiration, and plant C/N (Fig.2). The addition of elephants had mixed effects (Fig. 2). The sensitivity of soil C estimates to N deposition, soil C respiration and plant recycling increased, relative to the no elephant scenario (Fig. 2), whereas the sensitivity of plant C stocks decreased for most model parameters. The sensitivity of NPP and NECB estimates increased for most parameters in the With elephant scenario compared to the No elephant scenario (Fig. 2). Furthermore, model estimates were sensitive to elephant N uptake rate, and elephant recycling (urinary and fecal release) rate.

**Fig. 2.**
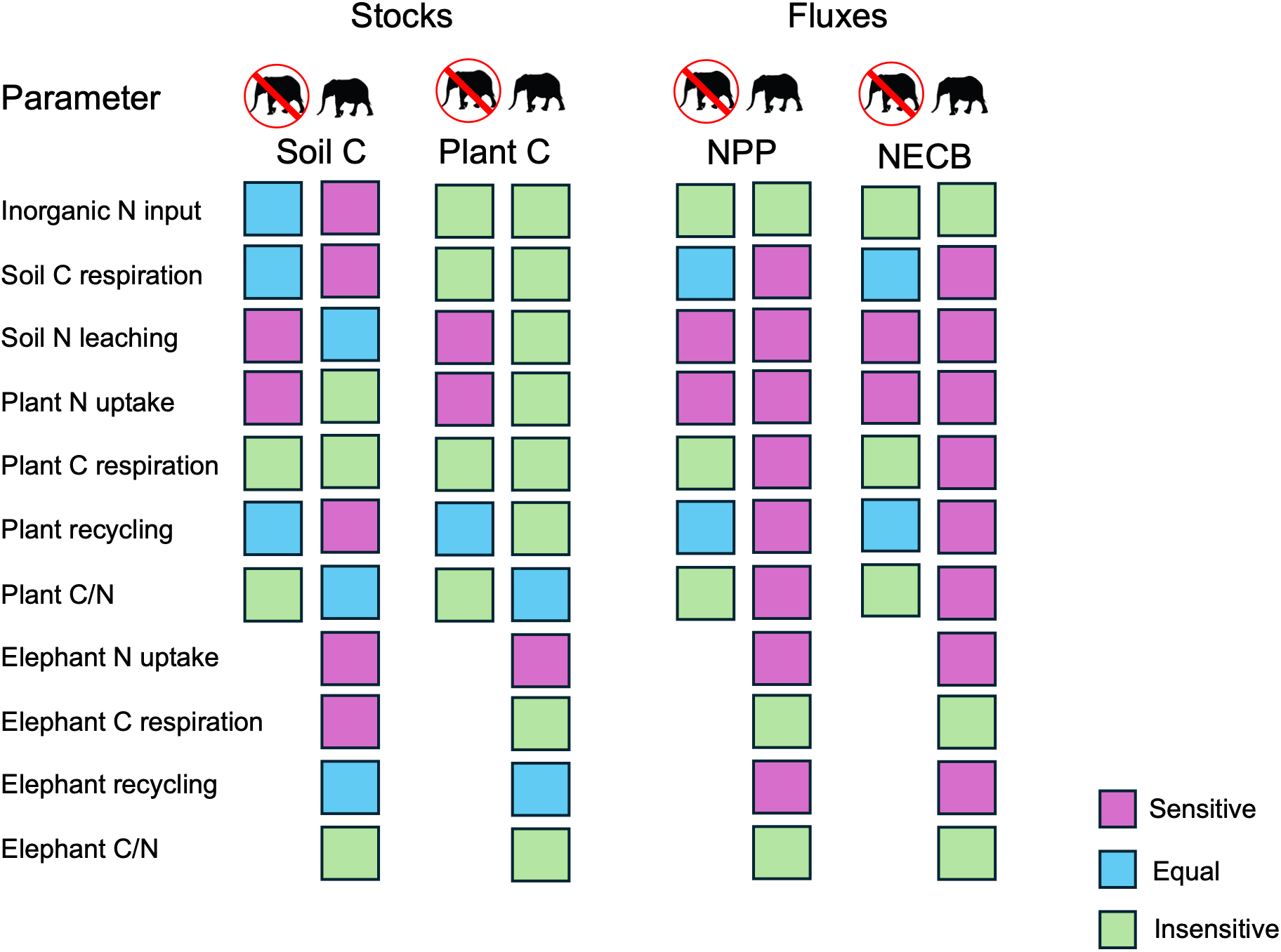
Sensitivity of carbon stock and flux estimates to ± 25% change in model parameter value presented in Table 2. The parameters were varied one-at-a time to assess sensitivity. A change in model estimate > 25% was considered *sensitive*, a change = 25% was considered *equal*, and a change < 25% was considered *insensitive*).

### 3.1 Estimates of the contribution of elephants to soil and plant carbon stocks

The values in Table 3 can be converted to t/ha—which are units generally considered in elephant conservation and management — by multiplying the Table 3 entries by a factor of 10x. The model estimates that mean soil C stock increases from 7.8 t/ha in the absence of elephants to 12.9 t/ha in the presence of elephants (a 1.7× increase). Thus, the elephant effects in the ecosystem could lead on average to an additional 5.1 t/ha of soil C, which would represent 40% of the average total soil C storage in the presence of elephants. However, the wide uncertainty range also indicates that elephant effects could cause soil stocks to decrease to 60% of that observed in their absence or increase by up to 5.3× more than in their absence (Fig. 3).

**Fig. 3.**
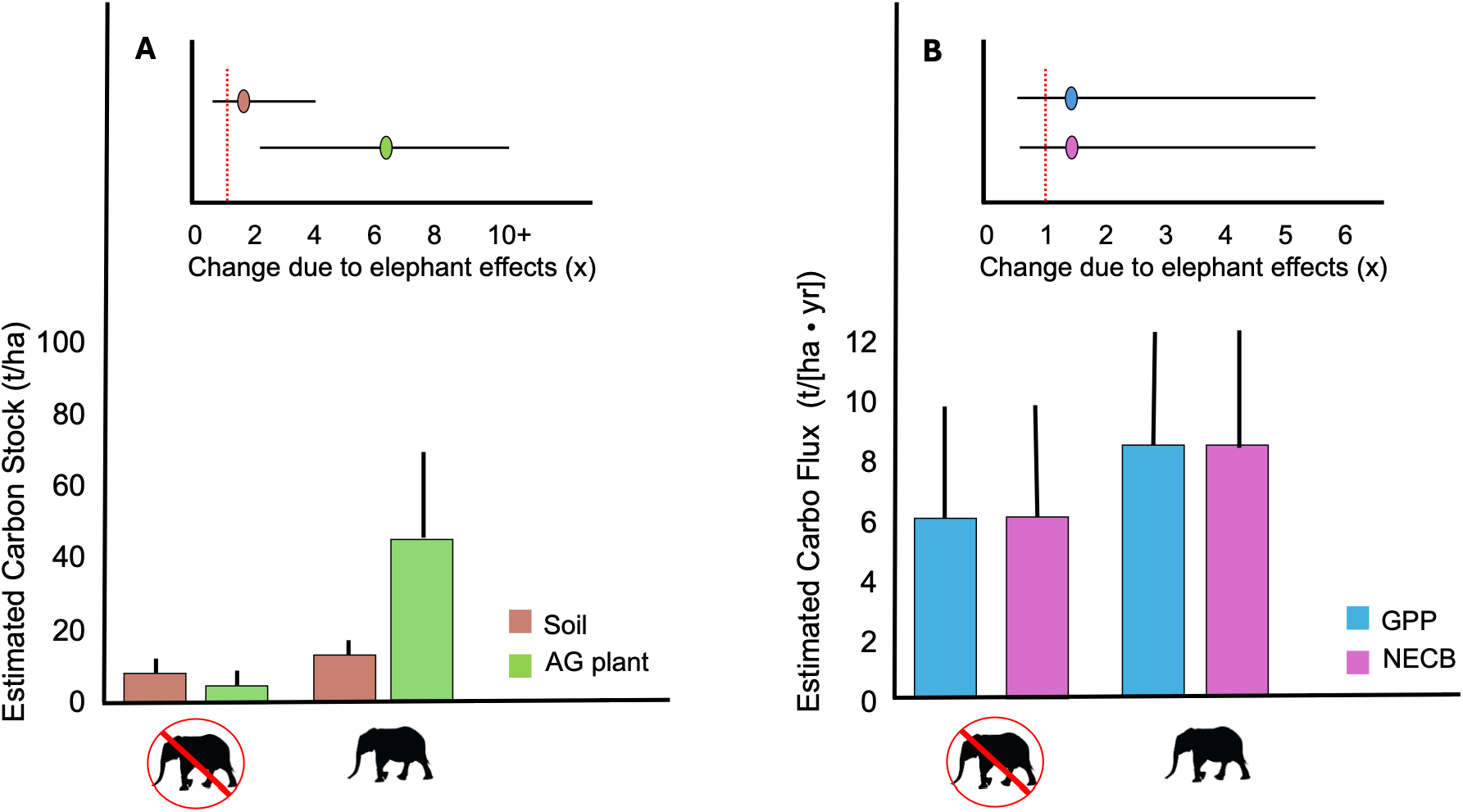
Mean and uncertainty_C-estimates_ ranges for model estimated soil and plant carbon stocks (A) and carbon fluxes (B) in the absence and presence of elephants. Insets in (A) and (B) present uncertainty ranges for elephant effects (uncertainty_EE_) on the calculated changes in carbon storage (stocks) or capture (fluxes) between the With elephant and No elephant scenarios.

Mean aboveground (AG) plant C stocks are estimated to increase from 7.2 t/ha in the absence of elephants to 47 t/ha in the presence of elephants (a 6.5 × increase), which amounts to a 39.8 t/ha increase (Fig. 3). This suggests that elephant effects could be responsible for 85% of the carbon stock in ecosystems with elephants present. However, there is a wide range of uncertainty, ranging from an approximate doubling of AG carbon storage to greater than 10 × increase.

### 3.2 Estimates of the contribution of elephants to carbon fluxes

The estimated daily NPP and NECB (Table 3) can be scaled to an annual basis by multiplying daily wet season fluxes by 181-days and dry season fluxes by 184-days and adding the two values (Fig. 3). Annual NECB accounts for NPP plus C losses due to soil respiration. The fact that the two estimates are equivalent in the No elephant scenario indicates that carbon losses due to soil respiration are negligible. Likewise, NPP and NECB are equivalent in the With elephant scenario. In this case, however, NECB accounts for additional C losses due to elephant respiration and enteric methane release. Again, the equivalence between NPP and NECB indicates that, at the modeled density of 0.5-0.7 animals per km^2^, carbon released directly from the elephants as CO_2_ and methane will have a negligible effect on the estimated ecosystem sink strength.

The model estimates that mean annual NPP and NECB could rise by 2.6 tC / (ha · yr) from a low of 6.1 tC / (ha · yr) in the absence of elephants to 8.7 tC / (ha · yr) in the presence of elephants, which is a 1.4× increase (Fig. 3). Elephant effects are estimated to be responsible for 30% of the estimated total carbon captured when they are present. However, the estimates carry a wide uncertainty range that includes a potential reduction in NPP and NECB, down to 40% of that estimated in the absence of elephants, or an increase by up to 5.5× more carbon captured in the presence of elephants.

## 4. Discussion

Nature-based climate solutions, which enlist natural ecosystem processes to capture atmospheric carbon and store it within ecosystems, are increasingly being promoted as important ways to help arrest and mitigate climate change (Griscom et al., 2020; Seddon et al., 2021). Accordingly, there is growing interest to consider savanna-woodlands as candidate ecosystems that could make important contributions to NbCS (Grace et al., 2006; Armani et al. 2022; Dobson et al. 2022; Zhou et al., 2023; Ossifo et al., 2026). However, recommendations for their conservation and management as NbCS focus on restoring and conserving vegetation and soils (Griscom et al., 2020; Zhou et al., 2023; Ossifo et al., 2026) and mostly overlook the functional role that animals may play. When animal effects are considered, such as the case of savanna elephants, it is often concluded that they may cause reductions in the capacity of savanna ecosystems to capture and store carbon, due to damages inflicted on aboveground vegetation (Pellegrini et al., 2017; Davies and Asner, 2019; Wigley et al., 2020; Wessels et al., 2026). Hence, savanna elephant conservation may be seen as antithetical to the goals NbCS. However, such conclusions are drawn from fragmentary considerations of the effects of savanna elephants on ecosystem carbon dynamics.

Here, we draw on general theory and modeling, as recommended by Jeltsch et al. 2000, to inform a more holistic exploration of some of the complexities of savanna ecosystem carbon dynamics resulting from elephant effects. The modeling, which accounts for elephant effects for an entire year including a 181-day wet season and a 184-day dry season, reveals that elephants could, on average, have a net positive effect on mopane ecosystem carbon capture and storage, despite consuming vegetation and felling trees. The uncertainty range of model estimates compare favorably with the range of empirical values reported in the scientific literature (Table 3), suggesting that the modeling is producing realistic estimates.

On average, estimates indicate that elephants could increase various components of net ecosystem carbon capture and storage by between 1.4-times to 6.5-times above that in their absence, which compares with other large grazers and browsers (Schmitz and Leroux, 2026). Thus, developing robust nature-positive climate solutions using savanna-woodland conservation and management (Zhou et al., 2023; Lisboa et al., 2025; Ossifo et al., 2026) will need to explicitly account for elephant effects to avoid producing inaccurate ecosystem carbon budgets.

The model estimates indicate that elephants could increase long-term steady state aboveground carbon stocks by 6.5-times above that estimated for the elephant absent scenario. This estimated increase does not align with results of elephant exclusion studies reporting that elephants significantly reduce aboveground biomass relative to no-elephant controls (Pellegrini et al., 2017; Wigley et al., 2020; Wessels et al., 2026). However, those studies report on reductions over short, decadal time frames. Longer-term studies and modeling of aboveground plant biomass in the presence of elephants (Ben-Shahar, 1996; Gordon et al., 2022) suggest that aboveground savanna vegetation biomass rebounds from initial declines over periods of 75-100 years, consistent with earlier hypotheses of the time-scale elephant effects on savanna-woodland vegetation (Caughley, 1976). Moreover a 6.5× increase in vegetation biomass in the presence of elephants compares with the magnitude of long-term (> 100 year) effects on aboveground vegetation measured and modeled for forest elephants in central Africa (Berzaghi et al. 2019).

The modeling further accounts for elephant effects on carbon stored in soils as organic matter, consistent with a holistic conceptualization of elephant effects on savanna-woodland ecosystem biogeochemical cycling (Skarpe et al., 2004; du Toit et al., 2014, Coetsee et al., 2019; Huntley, 2023). However, this particular aspect of elephant-driven carbon dynamics remains understudied (Hyvarinen et al., 2021; Sandhage-Hoffman et al., 2021). The available measures of elephant effects (Kinderman et al. 2025), corroborate model estimates that elephants can cause an average 1.7× increase in soil organic C stocks (Table 3). This increase could be enough to offset similar magnitudes of short-term aboveground biomass carbon losses (c.f., Wessels et al. 2026), especially because elephants may trample felled woody debris into soil storage pools (Sandhage- Hofmann et al., 2021; 2022). Thus, conclusions about the consequences of elephant conservation on NbCS based solely on aboveground effects will be inaccurate.

Because elephant net effects on ecosystem carbon stocks become manifest over long time periods (decades to centuries) it becomes challenging to verify whether elephant conservation is indeed enhancing NbCS. However, focusing measurement on carbon stocks overlooks measurements of fluxes (NPP and NECB), which are more responsive to short-term (interannual) changes in biophysical conditions (Veenendaal et al., 2004; Kutsch et al., 2008; Jin et al., 2013) and thus provide early (e.g., 1-5 year) indicators of ecosystem responses. Such measurements are currently not undertaken to assess elephant effects. The modeling suggests they should be measured because elephants could increase average annual NPP and NECB by 1.4×. This rise in productivity also explains why AG plant biomass carbon recovers over the long term. Analyses of fluxes are also necessary to assess the effects of land use and land cover changes (LULUCC) on carbon budgets of local savanna-woodland ecosystems to evaluate whether land conservation and management activities will avoid further emissions, let alone improve carbon sink strength (Ossifo et al., 2026). Accounting of emissions under LULUCC is also required as part of nationally determined contributions (NDC’s) under the UNFCCC biannual transparency reporting requirements.

The restoration and conservation of savanna-woodland ecosystems that includes the recovery of wildlife populations, especially elephants, is becoming an increasing priority throughout sub-Saharan Africa (van Aarde et al., 2006; Sandhage-Hoffman et al., 2022). Although efforts are in place to conserve large landscapes to facilitate elephant movement and prevent excessive local damage arising from high elephant densities within confined areas (Roever et al., 2013; Lindsay et al., 2017; Giliba et al., 2023; Simba et al., 2024), conservation efforts nonetheless continue to be anchored around existing protected conservation areas and wildlife game reserves (Roever et al., 2013; van Aarde et al., 2006; Giliba et al., 2023). Consequently, there is interest in managing elephant densities such that they are compatible with sustaining the vegetation structure within reserves and conservancies (Skarpe et al., 2004; Scheiter and Higgins, 2012; Thel et al., 2025). Suggested densities to achieve such goals range between 0.5-0.7 elephants per km^2^ (Scheiter and Higgins, 2012; Guldemond et al., 2017; Kuiper et al., 2026). However, the modeling presented here begs the further question: can those elephant densities lead to nature positive contributions to NbCS as well, and if so, by how much?

### 4.1 Case study of NbCS potential in a wildlife reserve

We examine the potential for elephant conservation within a wildlife reserve to contribute to NbCS using a case study of the 1958 km^2^ Chewore South Safari Area (CSSA) in northern Zimbabwe’s Zambezi Valley. The CSSA contains a diversity of habitats/ecosystem types, including riparian woodlands, miombo forests, open grasslands, and mopane savannas (Fig. 4), which support many wildlife and plant species (Tagwireyi and Chikwara 2025). These various habitats/ecosystem types are managed to promote biodiversity conservation, tourism, and ecosystem services that benefit local communities and help contribute to meeting global conservation goals. The region has a hot, wet season from November to April (∼181 days), with rainfall ranging between 600 and 900 mm and a cool, dry season from May to October (∼184 days) (Tagwireyi and Chikwara 2025).

**Fig. 4.**
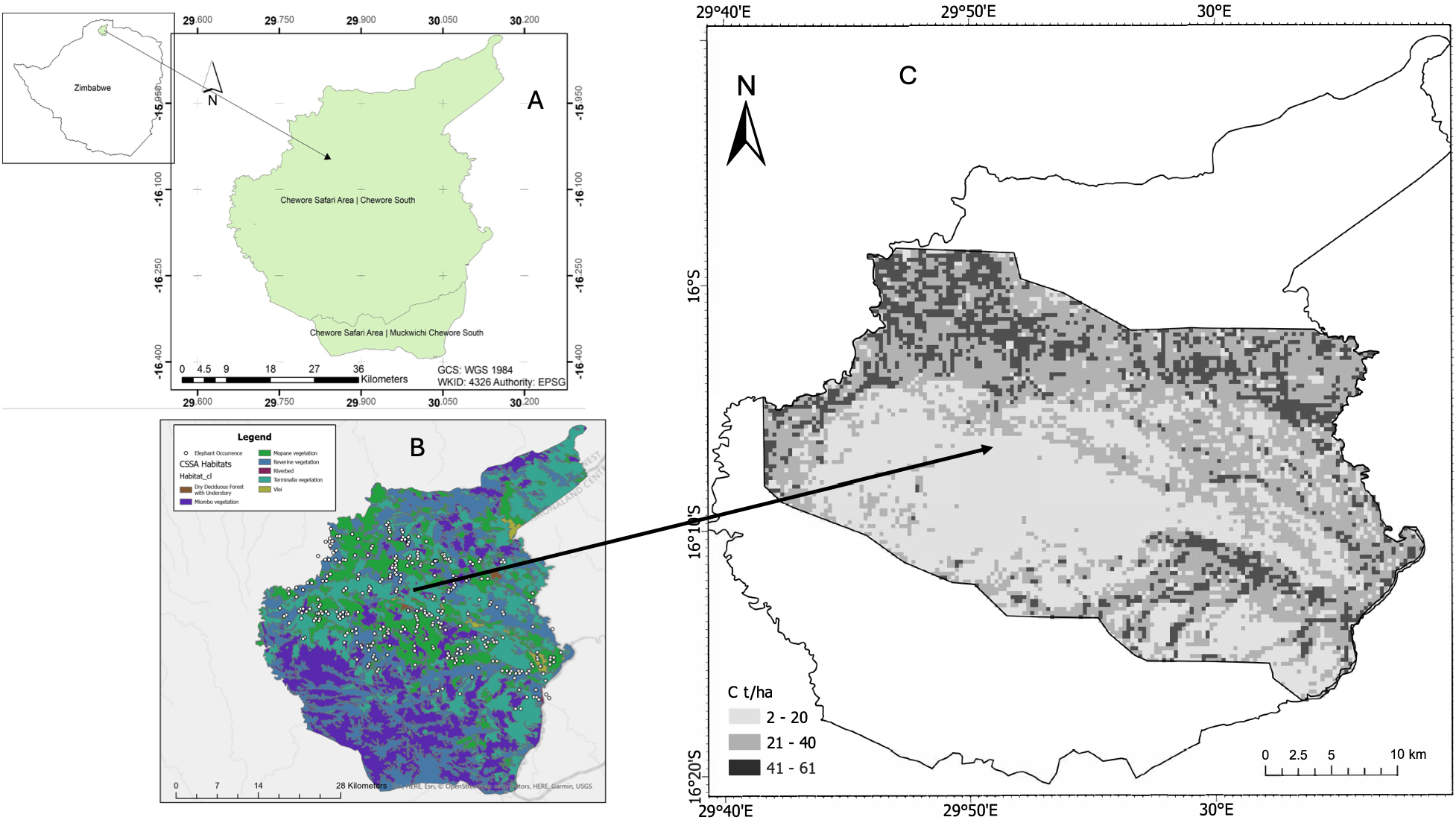
Maps depicting location and spatial occurrences of savanna elephants within mopane savanna portion the focal Chewore South Safari Area (CSSA) and aboveground carbon in mopane savanna. (A) Location of the CSSA within Zimbabwe. (B) Distribution of elephant occurrences measured in a 2025 aerial survey overlayed onto a landcover map depicting the spatial distribution of different habitat/ecosystem types. (C) Spatial distribution of aboveground carbon estimated using Google Earth Engine.

The savanna elephant is of conservation concern in the region owing to its ecological impacts and management significance (Tagwireyi et al. 2025). A management goal is to maintain elephant densities at approximately 0.3-0.5 individuals per km^2^, to avoid potential habitat destruction associated with higher elephant densities (Tagwireyi et al. 2025). An aerial survey conducted in September 2025 revealed that elephants within CSSA (Fig. 4) occurred predominately in the mopane and terminalia ecosystem types dominated by shrubs and trees of *Terminalia* and *Colophospermum mopane* (Fig. 4),.

We estimated aboveground vegetation biomass (AGB) using Google Earth Engine by combining spaceborne reference data with multitemporal imagery to estimate median aboveground biomass density, in t ha⁻¹, over CSSA by sampling 1,000 random points at 100 m resolution (see Appendix A: A3). The above-ground carbon density was then computed by multiplying each pixel’s predicted AGB by 0.47, the default carbon fraction for vegetation biomass (Appendix A: A3). The range in estimates of aboveground carbon stocks within this focal area (2.1 – 60.1 t/ha: Fig. 4) bracket the range of model estimated elephant effects on AG carbon (Table 3), suggesting that the range of model estimates of elephant-driven aboveground carbon are realistic. Assuming that model-estimated fluxes are equally realistic, then potential elephant-driven mean total carbon sink strength of the wildlife reserve by multiplying the estimated NECB values (Table 3) in t/ha by the area of the mopane-terminalia ecosystem within the CSSA landscape (which together cover 920.62 km^2^ = 9.2 x 10^4^ ha [Tagwireyi and Chikwara (2025)]). This translates into an estimated mean carbon uptake of 8 x 10^5^ tC/yr, or an estimated 2.9 million tonnes of CO2e captured annually in the area within the CSSA where elephants are present.

The estimated annual NECB for the intact ecosystem that includes elephants can be compared with emissions reporting for LULUCF in Zimbabwe’s 2025 Nationally Determined Contributions (NDC) Update Report to the UNFCCC (Government of Zimbabwe 2025: see table 4 in the Report). The estimated mean NECB is equivalent to 6% of reported national annual emissions from the land use and land use change and forestry (LULUCF) sector in 2025. It is equivalent to 7-10% of the annual emissions reduction goals between 2025 and 2030 in the LULUCF sector (Government of Zimbabwe 2025: see table 5 in the Report). The LULUCF sector identifies increasing area of natural forests, increasing area of planted forest, and reducing the extent of natural forests burned, and extent of wooded grasslands burned as top mitigation strategies. While this focus on restoring forests and protecting forests and grasslands from fire is an important way to leverage nature to mitigate climate change, the report does not identify other potential nature-based climate solutions among the proposed mitigation strategies. The evidence presented here reveals that the elephant effects at the landscape scale are sufficiently large that elephant conservation within the CSSA could be a promising NbCS. It thus argues for seriously considering and empirically quantifying elephant effects in mopane ecosystems within nature reserves and conservancies in Zimbabwe and throughout southern Africa more broadly, as a potential way to expand the portfolio of nature-based reductions of annual CO2e emissions.

### 4.2 Moving forward scientifically

The modeling presented here provides a critical first step in showing the feasibility of considering ACC via elephant conservation as part of a broader portfolio of NbCS to help arrest climate change and biodiversity loss together. The magnitudes of potential elephant effects argue for further investment in detailed measurements and refined modeling to improve the accuracy of carbon budget estimates, especially to address the wide range in uncertainty of the current estimates (Table 3; Fig. 3). An important value of the model, therefore, is that it strategically identifies what further work is needed to improve the accuracy of estimates of elephant effects on savanna ecosystem carbon budgets.

The uncertainty of model estimates hinge on several key parameters related to elephant impacts that are customarily not routinely measured in wildlife ecology nor in ecosystem science. The modeling reveals that the estimates of carbon capture and storage can be sensitive to the (in)accuracy of these parameter estimates:

a. *Elephant bioenergetics*. These include measures of elephant forage consumption rates in units of biomass per unit time, which can vary with forage availability and measures of dung and urine release in units of biomass mass per time under different forage and biophysical conditions (Appendix A: A2).
b. *Elephant effects on plant physiology*. Herbivory can cause reductions in plant photosynthesis and increases in plant respiration (Hawkes and Sullivan, 2001; Skarpe and Hester, 2008), which alters carbon captured per unit time and fixed as plant biomass (net primary productivity). Herbivory can also lead to plant reallocation of carbon from aboveground to belowground tissues (Skarpe and Hester 2008), which would question the use of aboveground carbon as the sole measure of plant response to elephant herbivory. More quantification of elephant effects on aboveground and belowground plant biomass is in order.
c. *Elephant ecosystem effects*. Elephants can impact ecosystem processes via trampling and waste release that alters soil respiration (microbial action) and nutrient (e.g., soil nitrogen) leaching losses, and the amounts of plant litter biomass released (plant recycling rate) to soils (Skarpe and Hester, 2008). More quantification of elephant effects on these aspects of ecosystem processes is needed.

There is a need to overcome an important challenge with the field research. While carbon storage in AG and soils can be well measured across different elephant densities (e.g., Ben-Shahar 1996, Woollen et al. 2012, Cambule et al. 2014, Sandhage-Hoffman et al. 2022, Kahumba and Tefera 2024, Kinderman et al. 2025), comparisons with locations that have similar biophysical conditions but with elephants absent (i.e., a control for elephant effects) are limited in number (Pellegrini et al., 2017; Wigley et al., 2020; Wessels et al., 2026). Without such study conditions it will remain challenging to validate empirically how much any gain in ecosystem carbon capture and storage can be attributed to elephant effects.

Finally, the model structure represents a simplification of complexity in savanna ecosystem structure. It treats plant vegetation as a single trophic compartment. More realistic models should consider the interplay between trees, shrubs and grasses (Fox et al., 2017; Ferry et al., 2021) given that interactions between these functional groups can determining amounts of plant and soil carbon stored (Zhou et al., 2023). Also, elephants are part of a community of many vertebrate herbivore species with myriad functional roles, interdependencies and feedbacks (Skarpe et al., 2004; Owen-Smith et al. 2019). Thus, future modeling may need to account for different herbivore functional roles (e.g., grazer, browser, mixed feeders) and their impacts on trees, shrubs and grasses. However, some evidence suggests elephant ecosystem effects may overshadow effects of other herbivores (Skarpe et al., 2004; 2014; Coverdale et al., 2016) suggesting our focus solely on elephant effects may provide a reasonable first approximation of animal-driven savanna ecosystem carbon cycling.

### 4.3 Summary

This report addresses an important knowledge gap about the feasibility of using savanna elephant conservation as a nature-based climate solution. The modeling provides a framework to help think holistically about the effects that elephants can have on savanna ecosystems and the potential levels of impact they may have on ecosystem carbon budgets. The modeling produces first approximation estimates that reveal the promise of savanna elephants to significantly enhance savanna ecosystem carbon capture and storage. The findings thus encourage investment to undertake further detailed field measurements and refined modeling to improve the accuracy of carbon budget estimates by reducing the uncertainty in field measurements that impact the model estimates, especially aboveground carbon storage. Such improved budget estimates have implications for improving the accuracy of NDC accounting among southern African countries.

## CRediT authorship contribution statement

OJS: Conceptualization, funding acquisition, formal analysis writing – original draft, reviewing and editing.

PT: Formal analysis, resources writing – reviewing and editing.

## Acknowledgements

This research was supported by funds from the International Fund for Animal Welfare.

## Appendix A

This appendix provides detailed information on the equilibrium solutions for the mathematical model used to estimate mopane ecosystem’s carbon budget in the absence and presence of savannah elephants. It is divided into 2. Section **A1** presents the equilibrium solutions for soil and plant carbon storage and carbon capture (NPP and NECB). Section **A2** presents a detailed explanation of the calculations used to estimate the model parameters. Section **A3** presents the methods used to quantify aboveground carbon storage for the case study of mopane ecosystem carbon capture and storage in the South Chewore Safari Area conservancy in northern Zimbabwe.

## A1. The model’s equilibrium solutions

The ecosystem model version used for the analysis (main text table 1) is a food chain comprised of three compartments: soil, mopane savanna plants, and elephants. The model tracks nitrogen and carbon inputs to the ecosystem (e.g., inorganic inputs, nutrient uptake, gross plant carbon uptake), outputs from the ecosystem (e.g., nitrogen leaching, and soil, plant and elephant respiration), and nitrogen and carbon recycling and storage within the ecosystem. At the ecosystem level, the model estimates net primary productivity (NPP) and net ecosystem carbon balance (NECB). NECB captures the net accumulation of carbon in ecosystems after accounting for the direct and indirect effects of animals on ecosystem carbon cycling (see Rizzuto et al. 2023 for more details).

The equilibrium solutions (i.e., a balance ecosystem carbon budget) for the model presented in table 1 is presented below for each of the two modeling scenarios: (1) carbon uptake and storage in the absence of elephants; (2) carbon uptake and storage in the presence of elephants.

### Soil and plant carbon stocks

The set of equilibrium solutions to estimate soil and plant elemental stocks for the scenario without elephants, derived by Rizzuto et al. 2023, is:

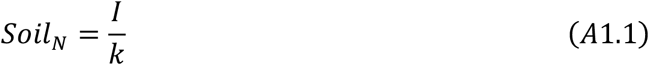

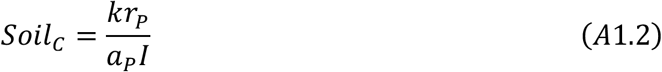

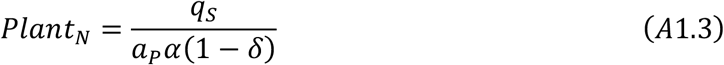

where N represents nitrogen and C represents carbon. The description of parameters is presented in the main text table 1. The model described in main text table 1, being stoichiometrically explicit, uses C:N ratios of plants (*α*) to track the budget and cycling of C in the ecosystem.

Thus, steady state plant C content is obtained as:

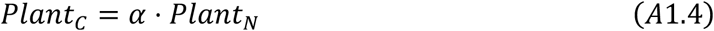

The set of equilibrium solutions for the scenario with elephants from Rizzuto et al. 2023 is:

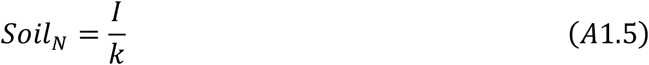

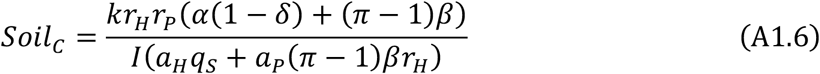

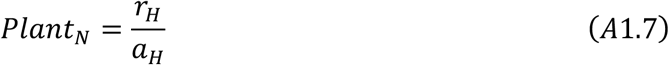

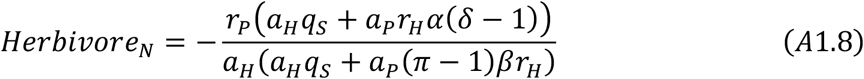

Again, being stoichiometrically explicit, the modeling uses C:N ratios of plants and elephants (*β*) to track the budget and cycling of C in the ecosystem. Thus, steady state plant C content is obtained as:

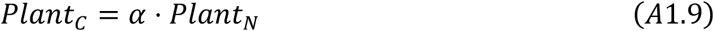

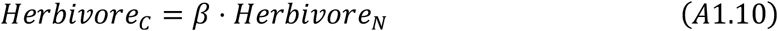

### Carbon capture

Net Ecosystem Carbon Balance (NECB) measures ecosystem carbon storage in terms of the net difference between an ecosystem’s anabolic and catabolic processes, i.e., the balance between net rate of carbon accumulation in ecosystems due to carbon fixation by plants (primary productivity), heterotrophic (i.e., animals, microbes) production and respiration, as well as additional losses including methane emissions directly from animals and soils and sediments of ecosystems (Chapin et al. 2006). We use equation A1.11 (see Rizzuto et al. 2023) to integrate the effects of plants and heterotrophs (animals and soil biota) in the accounting of NECB. Broadly, we define NECB as the sum of Net Primary Production and Net Heterotrophic Production (*NHP*):

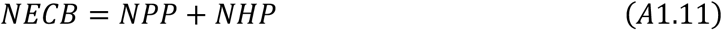

where *NPP* = *dP_C_/dt* (table 1). Equation A1.11 allows for debiting heterotrophic respiration from gross heterotrophic production to produce *NHP*. Thus, the *NPP* and *NHP* together capture the combined anabolic and catabolic processes happening in an ecosystem, across all trophic compartments. Net Primary Production is the balance of the photosynthetic and respiratory processes that happen in the autotroph (i.e., plants) compartment of an ecosystem, where:

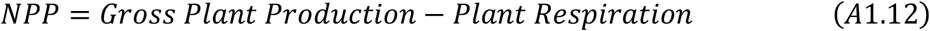

Additionally, Net Heterotrophic Production is the sum of all biomass-producing and respiratory processes taking place in the heterotrophic compartments of the ecosystem. These include (i) the herbivore (elephant) trophic compartment and (ii) the soil trophic compartment into which carbon from plant and animal waste and dead biomass is input, decomposed and recycled for nutrient uptake by plants (*N*) or stored (*C*). These considerations lead to the *NHP* estimate (Rizzuto et al. 2024):

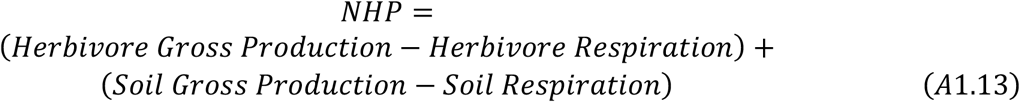

where Gross Production is measured as the C uptake rate by a given trophic compartment. Accounting for both components of heterotrophic C effects allows us to explicitly estimate the relative impact of different kinds of heterotrophs on NECB. Thus, for the elephant absent scenario (i), NECB is calculated as,

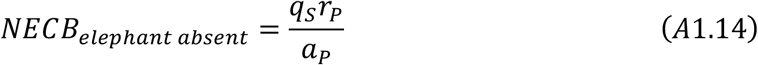

And, with elephant present scenario (ii), NECB is,

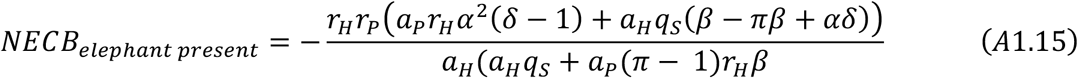

## A2. Parameters for the carbon cycle model

The estimated parameter values for the model were based on data obtained from the published scientific literature. The mean parameter values are presented in Table 2. The following narrative presents an explanation of how parameter values were calculated (including calculating a range in values, whenever possible) and the literature sources from which data were drawn.

### A2.1 Inorganic N inputs: kg N · (m^2^ d)^-1^

Atmospheric deposition wet and dry deposition = 7 – 9 Kg N ha^-1^ y^-1^ (Gayly-Lacuax et al. 2016) = 1.9 x10^-6^ – 2.5 x 10^-6^ kg N · (m^2^ d)^-1^

### A2.2 Soil C respiration rate: fraction of available soil C respired · (kg N m^2^ d)^-1^

The estimate of the parameter involved several steps to account for variation due to presence and absence of tree/shrub cover vs grass cover. The estimate required scaling per unit area respiration rate as a fraction of available soil carbon and then scaling that fraction to available soil nitrogen. This was done by prorating literature reported measures of soil respiration under trees/shrubs and under grass prorated by the relative abundance of tree/shrub and grass in a Mopane savanna.

Estimating available soil carbon required estimating soil carbon content per unit area, which is a product of literature reported values of soil % carbon in soils below tree/shrub vs grass and multiplying that by soil bulk density, prorated by the upper and lower range limit of soil bulk density measures in Mopane savanna. Soil nitrogen was likewise estimated as soil nitrogen content per unit area, which is a product of literature reported values of soil % nitrogen in soils below tree/shrub vs grass and multiplying that by soil bulk density, prorated by the upper and lower range limit of soil bulk density measures in Mopane savanna. The fraction of soil C respired in a 1 m^2^ area per day is then divided by the amount of soil nitrogen (in Kg) per m^2^.

The calculations are presented in the following.:

#### (i) Relative abundance of trees and shrubs, and grasses

Relative abundance of trees and shrubs per area = 0.29 (Feral et al. 2003) Relative abundance of grass per area = 0.71(Feral et al. 2003)

#### (ii) Available soil carbon

Soil organic carbon content - trees and shrubs =34.9 mg (g soil)^-1^ (Mlambo et al. 2007) = 0.035 g (g soil)^-1^

Soil organic carbon content - grasses =16.7 mg g soil^-1^ (Mlambo et al. 2007) = 0.017 g (g soil)^-1^

Soil bulk density = 1.6 -2 g soil cm^-3^ (Veenendaal et al. 2008). In a 10 cm soil core (Mlambo et al. 2007), soil mass becomes 16 – 20 g soil cm^-2^

*Prorated soil carbon content at lower bulk density* = (0.29 x 0.035 x 16) + (0.71 x 0.017 x 16) = 0.36 g C cm^-2^ = 3.6 x 10^3^ g C m^-2^

*Prorated soil carbon content at higher bulk density* = (0.29 x 0.034 x 20) + (0.71 x 0.017 x 20) = 0.44 g C cm^-2^ = 4.4 x 10^3^ g C m^-2^

#### (iii) Available soil nitrogen

Soil nitrogen content - trees and shrubs = 6.12 mg (g soil)^-1^ (Mlambo et al. 2007) = 0.0061g (g soil)^-1^

Soil nitrogen content - grass = 1.27 mg (g soil)^-1^ (Mlambo et al. 2007) = 0.0013 g (g soil)^-1^

*Prorated soil nitrogen content at lower bulk density* = (0.29 x 0.0061 x 16) + (0.71 x 0.0013 x 16) = 0.043 g N cm^-2^ = 4.3 x 10^2^ g N m^-2^ = 4.3 x 10^-1^ kg N m^-2^

*Prorated soil nitrogen content at higher bulk density* = (0.29 x 0.0061 x 20) + (0.71 x 0.0013 x 20) = 0.054 g N cm^-2^ = 5.4 x 10^2^ g N m^-2^ = 5.4 x 10^-1^ kg N m^-2^

#### (iv) *Soil respiration*

##### Wet season

Wet season average - trees and shrubs = 0.65 g CO_2_ m^-2^ hr^-1^ (Siele et al. 2008) = 4.18 g C m^-2^ d^-1^

Wet season average - grass = 0.60 g CO_2_ m^-2^ hr^-1^ (Siele et al. 2008) = 3.88 g C m^-2^ d^-1^

*Prorated respiration* = (0.29 x 4.18) + (0.71 x 3.88) = 3.97 g C m^-2^ d^-1^

#### Soil C respiration as fraction of available soil carbon

Upper = 3.97 g C m^-2^ d^-1^/ 3.6 x 10^3^ g C m^-2^ = 1.1 x 10^-3^ d^-1^ Lower = 3.97 g C m^-2^ d^-1^/ 4.4 x 10^3^ g C m^-2^ = 9.0 x 10^-4^ d^-1^

***Soil C respiration scaled to soil N* (fraction C · (kg N m^2^ d)^-1^)**

1.1 x 10^-3^ d^-1^ / 4.3 x 10^-1^ = 2.6 x 10^-3^

9.0 x 10^-4^ d^-1^ / 4.3 x 10^-1^= 2.1 x 10^-3^

1.1 x 10^-3^ d^-1^ / 5.4 x 10^-1^ = 2.0 x 10^-3^

9.0 x 10^-4^ d^-1^ / 5.4 x 10^-1^= 1.6 x 10^-3^

The modeling used the average value of 2.1 x 10^-3^ as a starting value.

#### Dry season

Dry season average - trees and shrubs = 0.12 g CO_2_ m^-2^ hr^-1^ (Siele et al. 2008) = 0.78 g C m^-2^ d^-1^ Dry season average - grass = 0.09 g CO_2_ m^-2^ hr^-1^ (Siele et al. 2008) = 0.58 g C m^-2^ d^-1^

*Prorated respiration* = (0.29 x 0.78) + (0.71 x 0.58) = 0.64 g C m^-2^ d^-1^

#### Soil C respiration as fraction of available soil carbon

Upper = 0.64 g C m^-2^ d^-1^/ 3.6 x 10^3^ g C m^-2^ = 1.7 x 10^-4^ d^-1^ Lower = 0.64 g C m^-2^ d^-1^/ 4.4 x 10^3^ g C m^-2^ = 1.4 x 10^-4^ d^-1^

***Soil C respiration scaled to soil N* (fraction C · (kg N m^2^ d)^-1^)**

1.7 x 10^-4^ d^-1^ / 4.28 x 10^-1^ = 3.9 x 10^-4^

1.4 x 10^-4^ d^-1^ / 4.28 x 10^-1^= 3.3 x 10^-4^

1.7 x 10^-4^ d^-1^ / 5.35 x 10^-1^ = 3.2 x 10^-4^

1.4 x 10^-4^ d^-1^ / 5.35 x 10^-1^= 2.6 x 10^-4^

The modeling used the average value of 2.9 x10^-4^ as a starting value.

#### (v) Elephant trampling effects

Elephant trampling can alter soil respiration. Data suggests that elephant effects can lead to between 0.1 x - 0.4 x reduction in soil respiration at low elephant densities (Sandhage-Hoffman et al., 2025).

The modeling assumed that respiration would conservatively be reduced by 0.1 x leading to 1.9 x10^-3^ as a starting value for the wet season, and 2.6 x10^-4^ for the dry season.

### A2.3 Soil N leaching rate: fraction · (m^2^ d)^-1^

Values for soil N leaching from savannas range between 1.9 × 10^-7^ - 1.9 × 10^-6^ (Aranibar et al. 2004, Coetsee et al. 2012). The modeling used an average 1.1 x 10^-6^ as a starting value.

Elephant presence and trampling can reduce leaching by 0.51 – 0.67 x (Maponga et al. 2022). Leaching rate was reduced by an average 0.55 x giving a starting value of 6.1 x 10^-7^.

### A2.4 Plant N uptake rate: fraction · d^-1^

Values for N uptake by savanna vegetation is on the order of 0.88 kg N ha^-1^ d^-^ (Abbadie et al. 1992, Cramer et al. 2012). Total available soil N has been estimated to be 100 kg N/ha (Cramer et al. 2012).

Thus, N uptake = 0.88 /100 = 8.8 x 10^-3^ for the wet season.

Dry season uptake = 0.6 x wet seasons uptake (Veenendaal et al. 2008), giving a value of 5.3 x 10^-3^.

While herbivory can limit the physiological regulation of plant N uptake, the magnitude of decline in N uptake remains unknown. Measures for the few studied terrestrial systems indicate that plant N uptake by grazed plants is 0.1 – 0.33 ∈ that of ungrazed plants (Zou et al. 2014, Guo et al. 2017). It was thus initially assumed that N uptake rate in the presence of herbivory is 0.24∈ that of uneaten plants.

Thus, plant N uptake = 2.4 x 10^-3^ with elephants in the wet season. Plant N uptake = 1.4 x 10^-3^ with elephants in the dry season.

### A2.5 Elephant N uptake dry and wet season: fraction N · (kg Elephant N d)^-1^

Elephant N uptake is estimated as % nitrogen in biomass consumed per day (2.2% in wet season; 1.1 % d^-1^ in dry season: Pretorius et al. 2012) divided by elephant body mass nitrogen (300 kg in a 4000 kg animal, i.e. 7.5% N: Reed et al. 2025) gives = 7.3 x 10^-3^ (wet season); = 3.7 x 10^-3^ (dry season).

However, elephant herds are comprised on average of between 43% and 70% non-adults (Leuthold 1976, Dunham 2024). Assuming non-adult elephants are on average 75% of the size of adults (Dunham 2024), give and estimated range in mean size of a herd of

High = (0.43 x 0.75 x 4000) + (0.53 x 4000) = 3410 kg

Low = (0.70 x 0.75 x 4000) + (0.30 x 4000) = 3300 kg

Or an average mass of 3355 kg.

Prorating consumption by average herd mass (i.e. 3355/4000 = 0.84) gives

6.1 x 10^-3^ (wet season); = 3.3 x 10^-3^ (dry season).

### A2.6 Plant C respiration rate: kg C· (m^2^ d)^-1^

Plant respiration is estimated to be 5.1 × 10^-3^ in wet season (Veenendaal et al. 2008)

Stress from herbivory is assumed to increase plant respiration leading to 3.9 x 10^-2^ for elephants in the wet season.

Plant respiration is reduced by 0.13x in dry season (Veenendahl et al. 2004). Thus dry season respiration is 6.6 × 10^-4^.

Respiration with elephants in the dry season is increased to 5.1 x 10^-3^.

### A2.7 Elephant respiration: kg C d^-1^

A 4000 kg animal is estimated to release = 4.65 kg C d^-1^ assuming DEE = 1.5 x BMR (Langman et al. 1995), and up to 11.77 kg C d^-1^ assuming calculated daily energy expenditure (DEE) of 3.8 x BMR (using estimating procedure from Schmitz et al. 2023, Box 1). Prorating for average elephant herd member body size gives 3.9 kg C d^-1^ assuming DEE = 1.5 x BMR; and up to 9.8 kg C d^-1^ assuming calculated daily energy expenditure (DEE) of 3.8 x BMR

### A2.8 Plant recycling rate: fraction · (kg N kg C m^2^ d)^-1^

#### Wet Season

Plant recycling rate is in the form of dead litter fall, which is estimated to be = 0.193 kg m^-2^ 181d^-1^ (Maponga et al. 2022). The comes from an estimated standing biomass of mopane vegetation = 20 Mg ha^-1^ (Colgan and Asner 2014) = 2 kg m^-2^. The average fraction of C in litter biomass = 513mg/g (Mlambo and Nyathi 2008) = 0.513 g/g. The average fraction N in litter biomass = 1.9% (Maponga et al. 2022) = 0.019 g/g.

However, empirical synthesis reveals that on average 45%-55% of annual litter produced in dryland ecosystems becomes fully decomposed within a year (Throop et al. 2025).

Thus, on a per m^2^ basis, a starting value of plant recycling rate = 0.5 x (0.193 kg m^-2^ y^-1^ / 2 kg m^-2^) / ((0.019 x 2 kg) · (0.513 x 2 kg) · m^2^ ·181 d)) = 7 x 10^-3^.

#### Dry season

Litter fall = 0.125 x wet season litterfall (Mlambo and Nyathi 2008) = 8.8 x 10^-4^

#### Elephant effects

Elephants can reduce amount of litter by 0.55 x (Maponga et al. 2022).

Thus, with elephants, litter fall in the wet season is reduced to 3.9 x 10^-3^

With elephants, litter fall in the dry season is reduced to 4.8 x 10^-4^ as a starting value.

### A2.9 Elephant recycling: fraction · d^-1^

#### Wet and Dry

##### Defecation

Defecation by an average 2500 kg forest elephant was calculated to be on average 43.8 kg/d dry mass in the wet season and 37.6 kg/d in the dry season (Ruggiero 1992). Assuming a 4000 kg average mass for adult savanna elephants (Owen-Smith 1988, Reed et al. 2025) and scaling up defecation by metabolic body mass (i.e., M^0.75^) ratios gives an estimated 60.4 kg/d in the wet season and 53.3 kg/d in the dry season. These values fall within the range of values measured by Coe (1972) for two adult individuals.

Prorating dung release for the average herd member mass of 3355 kg gives:

##### Wet season

60 kg/d x (3355/4000) = 50 kg/d

##### Dry season

53.3 kg/d x (3335/4000) = 45 kg/d

Assuming that elephant dung contains 0.55% N per unit mass (McCarthy et al. 2025), then 50 kg/d dung yields ∼0.28 kg N/d in the wet season and 45 kg/d yields ∼ 0.23 Kg N/d in the dry season. Scaling this as a percentage of elephant body mass N (where 4000 kg elephant contains 300 kg N, i.e., 7.5%N: Reed et al. 2025), gives an N release of an average herd as:

#### Wet season

0.28 kg N/d / (0.075 x 3355) = 1.1× 10^-3^/d of N

#### Dry season

0.23 kg N/d / (0.075 x 3355) = 0.96 × 10^-3^/d of N

#### Urine release

An average 4000 kg elephant releases 42-50 liters urine/d (Coe, 1972, Anathasubramanium 1979), or 42-50 kg urine/d. Multiplying these values by % N in urea (46.6%) and by percentage urea in urine (2%) gives a release of 0.39 – 0.47 kgN/d. Prorating for average herd mass gives a release of (3355/4000) x (0.39 – 0.47 kgN/d) = 0.32 – 0.39 kgN/d

Scaling these values as a fraction of elephant body mass N gives a release of 1.3 ×10^-3^ – 1.5 ×10^-3^/d.

#### Total release

Adding the values of urine release to values of defecation release gives an N recycling rate of: 2.4 ×10^-3^ /d - 2.6 ×10^-3^ /d for the wet season and 2.3 ×10^-3^ - 2.5 ×10^-3^ /d for the dry season.

### A2.10 Plant C/N

Estimate values range between 25.3 - 35.4 (Kahumba et al. 2024; Ries and Shugart 2008).

### A2.11 Elephant C/N

A 4000 kg elephant contains 2000 kg C and 300 kg N (Reed et al. 2025). Hence C/N = 6.67.

## A3. Methods used for the South Chewore Safari Area case study of carbon storage

We estimated aboveground carbon in the South Chewore Safari Area using data from satellite remote sensing. Following Zurqui et al. (2025) and Al Saim and Aly (2025), we estimated Aboveground Biomass (AGB) using Google Earth Engine (GEE) by combining spaceborne reference data with multitemporal imagery. In GEE we first assembled predictor layers by loading the Sentinel-2 Level-2A surface reflectance collection (COPERNICUS/S2_SR_HARMONIZED) and filtered it to 2023 - 2024. We masked clouds using the Sentinel-2 cloud-probability collection (COPERNICUS/S2_CLOUD_PROBABILITY) (keeping pixels with cloud probability <35% and excluding clouds/shadows flagged by the Scene Classification Layer). We then created median composites of the masked scenes and extracted bands B4 (red), B8 (near-IR) and B11 (SWIR). From these, we computed the Normalized Difference Vegetation Index (NDVI) and Normalized Difference Moisture Index (NDMI) as additional features. For radar data, we loaded the Sentinel-1 Ground Range Detected collection (COPERNICUS/S1_GRD) in Interferometric Wide (IW) mode for 2023 - 2024, selected the VV and VH polarization bands, and computed their median backscatter. We derived a new band VV–VH and applied a 50 m focal mean to smooth it. Elevation and slope were derived from the NASA SRTM 30 m DEM (USGS/SRTMGL1_003) using GEE’s ee. Terrain.slope function. We then obtained GEDI biomass reference data: the GEDI Level-4A raster product (LARSE/GEDI/GEDI04_A_002_MONTHLY). We then took the pixel-wise median of AGBD (aboveground biomass density, in Mg ha⁻¹ (aka t ha^-1^)over Chewore. We sampled up to 1,000 random points (seed = 42) from the stack of predictor images and the GEDI AGBD layer at 100 m resolution, discarding any samples with missing values.

Following Jiang et al. (2022) and Jiang et al. (2022) to model AGB, we trained a Random Forest (RF) regression using GEE’s SmileRandomForest classifier. The RF used 100 trees, 3 variables per split, minimum leaf population 5, bag fraction 0.6, and seed=42. The eight predictor variables (B4, B8, B11, NDVI, NDMI, VV–VH, elevation, slope) were regressed against the GEDI AGBD targets. For validation we performed a simple two-fold split by sample location (a non-spatial cross-validation) following Breiman, L. (2001). Finally, we applied the trained RF model wall-to-wall over Chewore to produce a 100 m map of AGBD.

